# Evidences of conditioned behavior in *Amoeba Proteus*

**DOI:** 10.1101/264176

**Authors:** Ildefonso M. De la Fuente, Carlos Bringas, Iker Malaina, María Fedetz, Alberto Pérez-Samartín, José I. López, Gorka Pérez-Yarza, María Dolores Boyano

## Abstract

Associative memory is the main type of learning wherein complex organisms endowed with evolved nervous systems respond efficiently to determined environmental stimuli. This fundamental cognitive property has been evidenced in different multicellular species, from cephalopods to Humans, but never in individual cells. Here, following Pavlov’s experiments with dogs that founded the principles of classical conditioning, we have observed the development of an associative memory in *Amoeba proteus,* which corresponds to the emergence of a new systemic motility pattern. In our cellular version of this conditioning behavior, we have used a controlled direct current electric field as the conditioned stimulus and a specific chemotactic peptide as the non-conditioned stimulus. Our study allowed us to demonstrate that *Amoeba proteus* are capable of linking two independent past events, and the induced associative memory can be recorded for up to at least four hours. For the first time, it has been observed that a systemic response to a specific stimulus can be modified by learning in unicellular organisms. This finding opens up a new framework in the understanding of the mechanisms underlying the complex systemic behavior involved in the cellular migration and the adaptive capacity of cells to the external medium.

---

One of the most remarkable accomplishments in the field of neuroscience is the description of a cohesive set of essential principles that define the nature of the basic forms of associative memory. Associative learning occurs through a process by which the connection between two previously unrelated stimuli, or a behavior and a stimulus, is learned; when such process befalls, it is assumed that the association of these stimuli is stored in a memory system.

For centuries, different thinkers have shaped a very plentiful and venerable history of research on basic learning processes. The combined work of philosophers, naturalists, physiologists, and life scientists has set the foundation upon which the modern learning theory currently stands. The most basic type of associative learning is the classical conditioning developed by the Laureate Nobel Price Ivan Pavlov, who established the first systematic study of basic principles of associative memory. In his studies, after appropriate conditioning, the dogs deprived of food were able to exhibit a consequent response –salivation-when a bell rung^1^.

Associative conditioning is ubiquitous in complex organisms endowed with evolved nervous systems, including all major vertebrate taxa and several invertebrate species. This complex process can also be reproduced and analyzed in artificial neural networks and different computational models^2^. Conditioned learning gives organisms the ability to adapt to ever-changing environments, and is considered a critical part of life’s regulation and survival. Despite its importance, associative memory has never been observed in individual cells.

Here, in order to determine whether conditioned behaviors are directly involved in cell migration, we analyzed the movement trajectories of *Amoeba proteus* under two external stimuli. Cellular migration is a critical systemic property of most cells, in fact, cellular life would be impossible without regulated motility. However, although some progress is being made in understanding this process, how cells move efficiently through diverse environments, and migrate in the presence of complex cues, is an important unresolved issue in contemporary Biology. Free cells need to regulate their locomotion movements in order to accomplish critical activities like locating food and avoiding predators or adverse conditions. In the same way, cellular migration is required in multicellular organisms for a plethora of fundamental physiological processes such as embryogenesis, organogenesis and immune responses. Indeed, deregulated human cellular migration is involved in important diseases such as immunodeficiencies and cancer^3, 4^.

In our study of cellular motility in amoebae, we have used an appropriate electric field as conditioned stimulus and a specific peptide as chemo-attractant. Previous experimental studies have shown that *Amoeba proteus* exhibit galvanotaxis, that is, a directed movement in response to an electric field. In fact, it has been described that practically 100% of the amoebae migrate towards the cathode for long periods of time under a direct current electric field (dcEF) in a range between 300 mV/mm and 600 mV/mm^5^. Likewise, amoebae are known to present chemotactic behaviors, in particular, the peptide nFMLP, typically secreted by bacteria, is able to provoke a strong chemotactic response in many different kinds of cells. The presence of this peptide in the environment may indicate the amoeba that food organisms might be nearby. In addition, nFMLP at specific concentrations is able to sharply increase the amoeba’s phagocytic behavior^6^.

Our experiments were carried out on a set-up that allowed us to simultaneously expose the amoeba to both stimuli (Figure 1). This system consists of two standard electrophoresis blocks, about 17.5 cm long, one directly plugged into a normal power supply and a second one connected to the first via two agar bridges that transfer the current from one block to the other while preventing the direct contact of both the anode and the cathode with the medium where the cells are located. In the middle of the second electrophoresis block we placed a glass structure that allowed us to create a laminar flux that not only permitted the electric current to pass through, but also generated an nFMLP peptide gradient that the amoebae were able to detect and respond to (see Methods section for more details). All the experiments were carried out in Chalkley’s medium, a standard, nutrient-free saline medium for *Amoeba proteus* at ambient temperature, 19-20°C.

**Figure 1.**
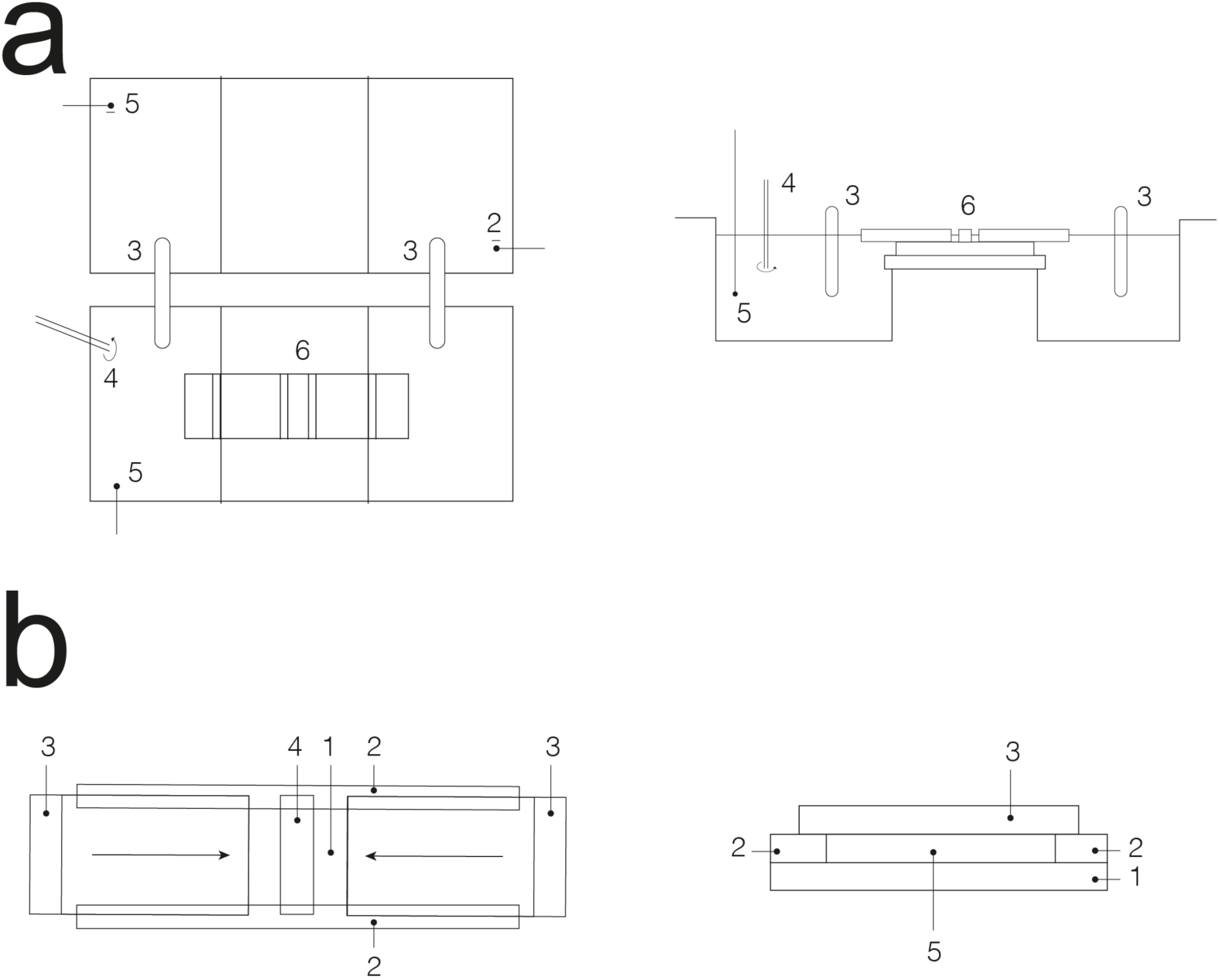
Experimental set-up. Panel a shows the top and lateral views of the experimental system. 1: anode; 2: cathode; 3: agar bridges, 8 cm long; 4: chemotactic peptide; 5: probe electrode used to monitor the electric field; 6: glass structure. Panel **b** corresponds to the top and frontal views of the glass structure in which the cells are placed. 1: standard slice glass 75 × 25 mm; 2: glued cover glasses 0.1 mm tall; 3: cover glasses that enclose the structure, each 4 cm long; 4: thin cover glass, about 3 mm wide under which the cells are placed; 5: experimental chamber 0.1-0.2 mm tall where the cells are subjected to the experiments.

1. Migration in the absence of stimuli. First, we have studied the locomotion movements of *Amoeba proteus* in the absence of stimuli. To this end, groups of 5-8 cells up to a total of 50 were placed in the middle of the glass set-up and their respective trajectories were recorded for 30 minutes. Figure 2a showed that the cells migrated in all directions. The directionality of each amoeba was quantified by the cosine of the displacement angle^5^; in our case, we considered the angle formed by the position after 30 minutes of displacement and the origin of the movement (see Methods). The results showed that the values ranged between -1 and 0.995, being -0.116 ± 1.56 the median ± Interquartile range (instead of mean ± SD, results were depicted as median ± IQ because the values were not normally distributed). These observations indicated that without the presence of any stimulus, amoebae moved in any direction randomly.
2. Directionality in electric field (galvanotaxis). The galvanotactic behavior of *Amoeba proteus* was analyzed in groups of 5 to 8 cells (up to a total of 50) subjected to a controlled electric field. The observations showed that 100% of the amoeba migrated towards the cathode (Figure 2b), matching with previously reported results^5^. The values of the cosines of displacements were distributed between 0.037 and 0.999 (0.994 ± 0.03 median ± IQ). In order to quantify the significance of our results, we performed a non-parametric test (Wilcoxon ranksum test) to compare the distributions of the cosines of the displacement in both situations, without stimulus and under the presence of the electric field. The p-value of the test was 10^-13^, which rejected the hypothesis that both samples came from the same distribution, or in other words, the result indicated that the behaviors with and without the stimulus of the electric field were significantly different.
3. Chemotactic gradient (chemotaxis). The behavior of amoebae during chemotaxis was analyzed by exposing the cells (50 in total) to an nFMLP peptide gradient placed in the left side of the setup. Figure 2c shows that 86% of the cells migrated towards the chemotactic gradient (left). The cosines of the displacement angles ranged between -0.997 and 0.987 (-0.825 ± 0.72 median ± IQ). Then we performed two comparisons; first, between the cosines obtained from the experiment with the chemotactic gradient and without the presence of the stimulus (p-value = 10^-4^), and second, between the cosines under chemotactic gradient and with the presence of electric field (p-value = 10^-17^). These tests indicated that the behavior under the chemotactic gradient was significantly different to both, the absence of stimulus, and the presence of electric field.
4. Double stimulus (cellular induction process). In this case, the motility of cells (up to 210 cells in groups of 5-8) was studied when subjected to both stimuli at the same time. We placed the cathode on the right of the set-up and the anode with the nFMLP peptide gradient in the left. The experiments were recorded for a total time of 30 minutes (Figure 2c). The results showed that 20.38% of the cells ignored the directionality of the electric field and moved towards the anode-peptide, favoring the chemotactic stimulus while the remaining cells migrated to the cathode (Figure 2d). The cosines of the displacement angles were distributed between -1 and 1 (0.978 ± 0.41 median ± IQ). The difference between the Interquartile ranges of this experiment and the experiment with only an electric field indicated that a new behavior had appeared among a group of cells. This new behavior was characterized by the migration towards the anode. Moreover, the statistical test comparing these two experiments corroborated this new behavior, with a p-value of 0.006.
5. Conditioned behavior test To study whether the migration towards the anode was persistent and could be reproduced in subsequent galvanotaxis experiments we manually extracted these amoebae that displayed the new behavior and placed them on normal culture medium in a small Petri dish in the absence of stimuli for about 5 minutes. Afterwards, the cells, usually in groups of one to three, were placed on a new identical glass and block set-up that had never been in contact with the chemotactic peptide nFMLP. Then, the amoebae were subjected once again to a single electric field, as described above. The results showed that 98% of the cells moved towards the anode where the chemotactic peptide was absent (Figure 2e). In this case, the cosines of displacements ranged between -1 and 0.104 (-0.997 ± 0.02, median ± IQ), indicating that the majority of cells moved in the direction of the anode, which corroborated that a new behavior had appeared among those cells. Finally, in order to calculate the persistence time of this new systemic motility pattern we placed the cells in nutrient-free Petri dishes in the absence of stimuli for periods of 30 minutes and exposed them once again to a controlled electric field (galvanotaxis). This process was repeated several times and allowed us to demonstrate that the new systemic motility pattern prevailed for up to at least 240 minutes in some *Amoeba proteus*.

**Figure 2.**
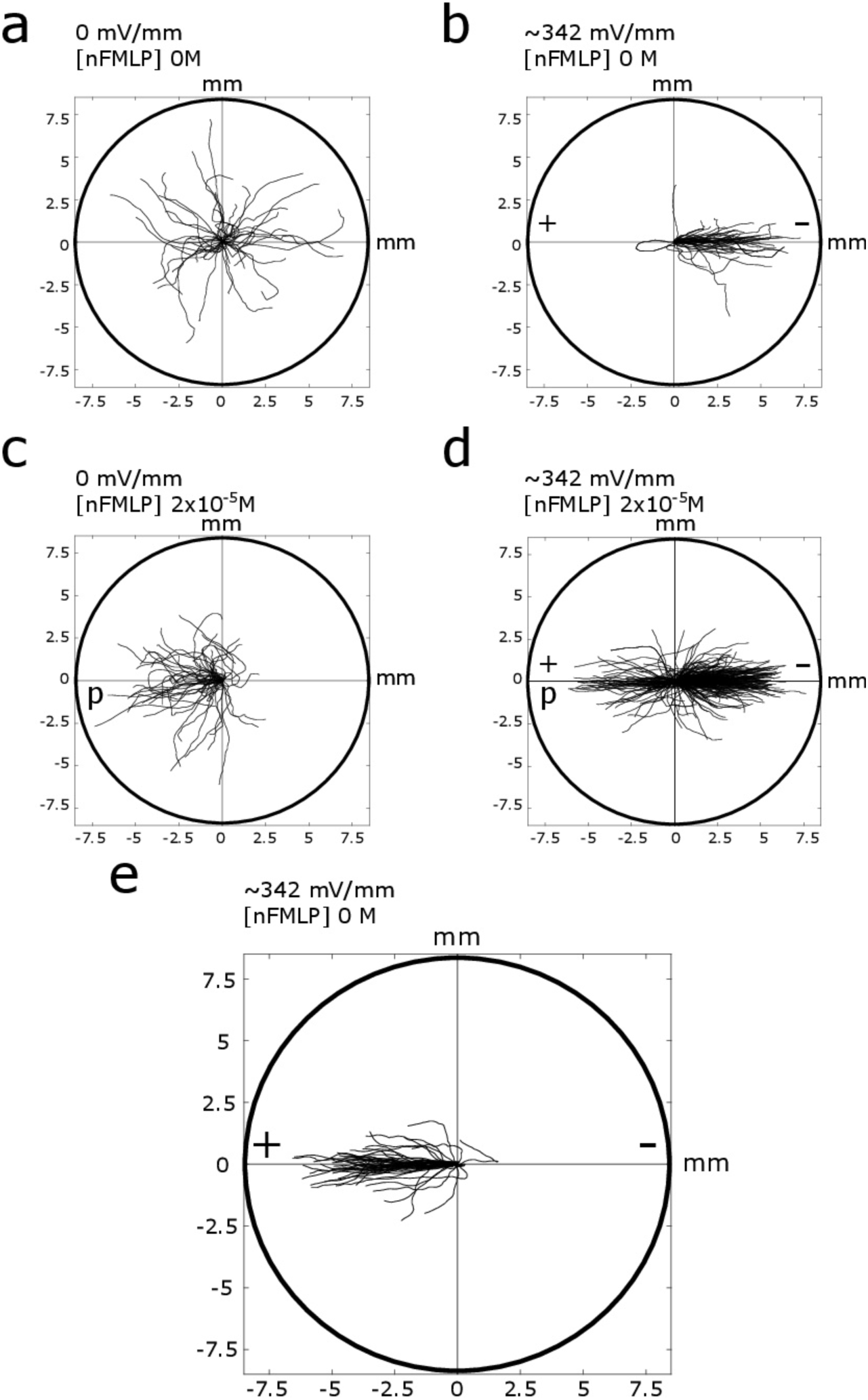
Migration trajectories. Composite trajectories of *Amoeba proteus*. The initial location of each cell has been placed at the center of the diagram. All the trajectories shown are 30 minutes long. The cathode of the electric field was always placed at the right, while the chemotactic peptide nFMLP and the anode were always placed at the left. **a,** shows the locomotion movements of amoebae in the absence of stimuli (n = 50). **b,** corresponds the response to a single electric field (n = 50). In panel **c,** we can observe the trajectories of amoebae (n = 50) subjected to a single chemotactic gradient. **d,** shows the trajectories of cells (n = 210) in response to a chemotactic gradient and electric field simultaneously (conditioning). **e,** corresponds to the migration movements of conditioned amoeba (n = 50), in which the cells develop a new motility pattern characterized by movements towards the anode in the absence of peptide. In all the diagrams, both the x and y axis show the distance in mm.

The current, generalized opinion is that cells are complex molecular genoteques (molecular boxes governed by genes) in constant evolution that lack the capacity to associate unrelated stimuli, record them and learn new systemic behaviors in order to adapt to the external medium in a flexible way.

Here, by using an appropriate direct current electric field (galvanotaxis) and a specific peptide (nFMLP) as a chemoattractant (chemotaxis) we have addressed essential aspects of the *Amoeba proteus* migration. We have found that cells were capable of linking two past events, shaping an associative memory that led to the emergence of a new systemic motility pattern. This induced associative memory prevailed in the cells for up to at least 240 minutes.

We have first observed that practically all the amoebae exhibited an unequivocal behavior characterized by the migration towards the cathode when exposed to an electric field under specific conditions (Figure 2b). However, if the amoebae were subjected to a simultaneous chemotactic and galvanotactic stimulus (induction) placing the peptide in the anode, some cells that we call induced cells (about 20% of the total) did migrate towards the peptide and were able to acquire a new and singular systemic response in their cellular locomotion (Figure 2d). This new cell behavior could be reproduced in subsequent galvanotaxis experiments and consisted on the migration towards the anode in the absence of the peptide (Figure 2d).

It must be remembered that all the cells migrated towards the cathode before the induction. During the induction process only 20.38% of the amebae moved to the site where the peptide was located (anode). Subsequent experiments demonstrated that, in the absence of peptide in the anode, practically none of these cells migrated towards the cathode, as it would be expected, and instead, 98% of them still moved to the anode where no peptide was placed (Figure 2e). Thus, we can conclude that these induced cells seem to have acquired a new behavior of systemic motility after being exposed to an induction process consisting in two simultaneous stimuli for some time (30 minutes).

In other words, our experiments showed that when the exposition to a stimulus related to the amoeba’s nourishment (peptide) is accompanied by an electric field, and the peptide is placed in the anode, the amoeba appear to associate the anode to food (the peptide) because after 30 minutes of induction the amoebae developed a new pattern of cellular motility characterized by movements towards the anode even in the absence of nourishment (peptide) there.

The natural behavior of practically all amoeba is to go to the cathode, under specific galvanotactic conditions^5^, and it is yet unknown exactly why this migration pattern occurs. However, after a process of induction, the amoebae seem to associate food with the anode and, consequently, modify their conduct, behaving against their natural tendency to move to the cathode. Strikingly, this induced association of anode and food can be remembered for relatively long periods of time. In our experiments this newly learned systemic motility pattern prevailed for up to at least 240 minutes. It is necessary to note that the cellular cycle of *Amoeba proteus*, although it varies depending on the environment, is usually about 24 hours long under controlled culture conditions^7^.

In pavlovian terms, the electric field, specifically the anode, represents the “bell” (the conditioned stimulus) while the peptide would be the “food” (unconditioned stimulus) and the cellular migration corresponds to the dog’s “salivation” (unconditioned response). The peptide (unconditioned stimulus) attracts some of the cells, and since it is always placed in the positive pole of the electric field, the amoeba associates the anode with the peptide. After the conditioning (“induction”), both stimuli remain linked in the cell for a relatively long period of time. Consequently, the amoeba’s movement will respond later on to the presence of an electric field by migrating towards the anode. Pavlov called this process the “conditional response”.

In our cellular version of Pavlov’s experiments, we have studied a classical pavlovian conditioning called “simultaneous conditioning”, in which the conditioned and unconditioned stimulus are presented at the same time. Our results showed that practically all the conditioned *Amoeba proteus* exhibit the ability to learn and remember the relationship between the two stimuli, that is, the cells can “learn” via conditioning, similarly to the dogs in the classic Pavlov’s experiment. Noteworthy, the fact that individual cells are able to acquire learned associations to guide their complex migration movements has never been verified so far.

We do not know the molecular mechanisms by which this cellular associative memory is sustained. However, different studies at the cellular level suggest some metabolic processes that could be involved. For instance, Hopfield-like attractor dynamics have been observed in self-organized enzymatic networks which have the capacity to store functional catalytic patterns that can be correctly recovered by specific input stimuli^8^. In addition, evidences of functional memory which can be embedded in multiple stable molecular marks during epigenetic processes have been reported^9^. On the other hand, long-term correlations (mimicking short-term memory in neuronal systems) have also been analyzed in experimental calcium-activated chloride fluxes belonging to *Xenopus laevis* oocytes^10^. Likewise, similar correlations have been observed in other different cellular processes not related to the neuronal lineage10. Finally, the presence of non-trivial correlations in the hunting movements of enucleated *Amoeba proteus* has been recently verified^11^.

The mechanisms underlying cell motility are extremely complex and the ability of cells to direct their movement and growth in response to external stimuli is of critical significance for cell functionality. Cell motility has also extreme relevance on physiological processes and diseases in Humans such as organogenesis, morphogenesis, tissue repair and cancer metastases.

Here, for the first time, it has been observed that unicellular organisms are able to modify their systemic response to a determined stimulus implicated in motility exclusively by learning. This fact opens up a new framework in the understanding of the mechanisms that underlie the complex systemic behavior involved in cellular migration and in the adaptive capacity of cells to the external medium.

## Methods

### Cell Cultures

*Amoeba proteus* (Carolina Biological Supply Company, Burlington, NC. Item # 131306) were grown at 21°C on Simplified Chalkley’s Medium (NaCl, 1.4 mM; KCl, 0.26 mM; CaCl2, 0.01 mM), alongside *Chilomonas* as food organisms (Carolina Biological Supply Company Item #131734) and baked wheat corns.

### Experimental Set-up

The set-up (Figure 1) consisted mainly of two standard electrophoresis blocks, 17.5 cm long (Biorad Mini-Sub cell GT), a power supply (Biorad 3000Xi Computer Controlled Power Supply), two agar bridges (2% agar in 0.5n KCl, 8 cm long) and a structure made from a standard glass slide and covers (commonly used in cytology).

The first electrophoresis block was directly plugged into the power supply while the other was connected to the first block via the two agar bridges, which allowed the current to pass through and prevented the direct contact between the anode and cathode and the medium where the cells would be placed later. Both electrophoresis blocks consisted of 3 parts: on the extremes, there are 2 wells 5.6 cm deep which were filled by the conductive medium and, in the middle an elevated platform about 5 cm tall that separated them, see Figure 1a and b.

The glass structure was placed in the middle of the elevated platform of the second block. This structure consisted of two 60x24x0.1 cover glasses glued to the sides of a standard 75x25 mm glass slide by a thin layer of silicone. The cover glasses were placed in a way that shaped a rectangular chamber, see Figure 1c. To cover this structure, we used 3 different rectangular fragments of cover glass, a thin, usually about 3 mm wide glass that is placed in the middle and two approximately 4 cm long glasses to the sides.

### Cell preparation

All amoebae were previously placed in a Petri dish for about 24 hours in Chalkley’s medium on a plastic Petri dish and in the absence of external stimuli. Only healthy amoebae, strongly attached to the dish must be considered in the experiments.

The cells were washed in clean Chalkley’s medium and placed in the middle of the glass set-up in groups of 5 to 8, under the thin 3 mm cover glass (Figure 3c) and left to rest for 5 minutes or until all of them appeared to be firmly attached to the bottom of the experimental chamber. Next, the two 4 cm long cover glasses were placed on the sides of the glass structure, protruding outside of the middle platform of the block and over the lateral wells (figure 1a). After, each well was filled using 75 ml of Chalkley’s medium, in such a way that the glass protrusion over each well is in contact with the liquid’s surface. Finally, as the Chalkley’s medium slowly filled up the experimental chamber, both lateral cover glasses had to be gently pushed towards each other until they touched the middle cover glass, completely covering the whole structure and forming a laminar flux that connected both lateral wells.

### Electric field (galvanotaxis)

An electric field of constant 60 V was applied to the first electrophoresis block, which was then conducted to the second by the two agar bridges. Direct measurements taken with a multimeter in the second block showed that the strength of the electric current in the oscillated between 58.5 and 60 V (334-342 mV/mm) while the intensity values oscillated between 0.09 and 0.13 mA.

After 30 minutes of exposure, during which the cellular migration movement were recorded, the power supply was turned off and the agar bridges removed.

All the experiments where the only stimulus was an electric field were performed in an electrophoresis block that had never been in contact with any chemotactic substance.

### Peptide gradient (chemotaxis)

In the left well of block two, 75ul of 2x10^-4^M nFMLP (Sigma-Aldrich F3506) peptide solution was diluted for a final concentration of 2x10^-5^M. In order to homogenize the solution and accelerate the creation of a chemotactic gradient in the experimental chamber we carefully shook the left well for 2 minutes. Finally, the cells behavior was recorded for 30 minutes.

### Track recording and digitizing

The motility of the cells was recorded using a digital camera attached to a SM-2T stereomicroscope. Images were acquired every 60 seconds, over a period of about 30-33 minutes (30-33 frames). Since automated tracking software is often inaccurate^12^, we performed manual tracking using the TrackMate software in ImageJ (http://fiji.sc/TrackMate), as strongly suggested elsewhere^12^. Each track corresponds to an individual amoeba.

### Directionality analysis and statistical significance

In order to quantify and compare the directionality of cell migration towards the anode or the cathode, we computed the cosines of the angles of displacement of each amoeba^5^. More precisely, we calculated the cosine of the angle formed between the start and final positions of each cell. Consequently, we were able to study quantitatively if an amoeba moved towards the cathode (positive values), or towards the anode (negative values of the cosine). In addition, this study suggested the degree of directionality, since values closer to 1 (or to -1 in the case of the anode) indicated a very high preference towards that pole. Next, to estimate the significance of our results, first we studied if the distribution of cosines of angles came from a normal distribution, by applying the Kolmogorov-Smirnov test for single samples. Since the normality was rejected, the groups of cosines were compared in pairs by a non-parametric test, the Wilcoxon ranksum test.

## Acknowledgements

We would like to thank Laura Pérez Gomez for her assistance designing figure 1. This work was supported by a grant of the UPV/EHU, GIU17/066, and the Basque Government grant IT974-16.

## Author contributions

CB, IMDF: Designed experiments; CB, MF, MDB, APS, GPY, JIL, and IMDF: Performed the experiments; IM: Performed the quantitative analysis. All authors wrote the manuscript and agreed in its submission; IMDF: Conceived and directed the investigation.

